# Biosynthesis of the Multifunctional Isopropylstilbene in *Photorhabdus laumondii* Involves Cross-Talk between Specialized and Primary Metabolism

**DOI:** 10.1101/2022.10.20.513026

**Authors:** Siyar Kavakli, Gina L. C. Grammbitter, Helge B. Bode

## Abstract

Isopropylstilbene (IPS) derived from the entomopathogenic bacterium *Photorhabdus* represents the only known stilbene which is not produced by a plant stilbene synthase but a bacterial PKS II synthase. While the exclusive cyclization reaction, responsible for the formation of the characteristic iso-branched side-chain of the molecule, was studied in the past, some parts of the biosynthetic route remained elusive. In this study, we revealed the role of StlB that is able to produce CoA-derivatives and demonstrated the elongation of cinnamoyl-CoA with enzymes from the bacterial fatty acid biosynthesis pathway. Thus, we deciphered cross-talk between the enzymes from primary and specialized metabolism. These insights led, for the first time, to the production of IPS in a heterologous host.

## 1. Introduction

Stilbenes are well-known natural products usually produced by type III PKS systems (or stilbene synthases, STS) in plants. The biosynthetic mechanism is well-described and involves elongation of cinnamoyl- or coumaroyl-CoA thioesters with three malonyl-extender units.^1–3^ Subsequent cyclization and aromatization leads to the formation of stilbenes, giving rise to resveratrol, pterostilbene and pinosylvin as some examples for this compound class, which is widespread in plants (Fig. 1).^4^

**Figure 1.**
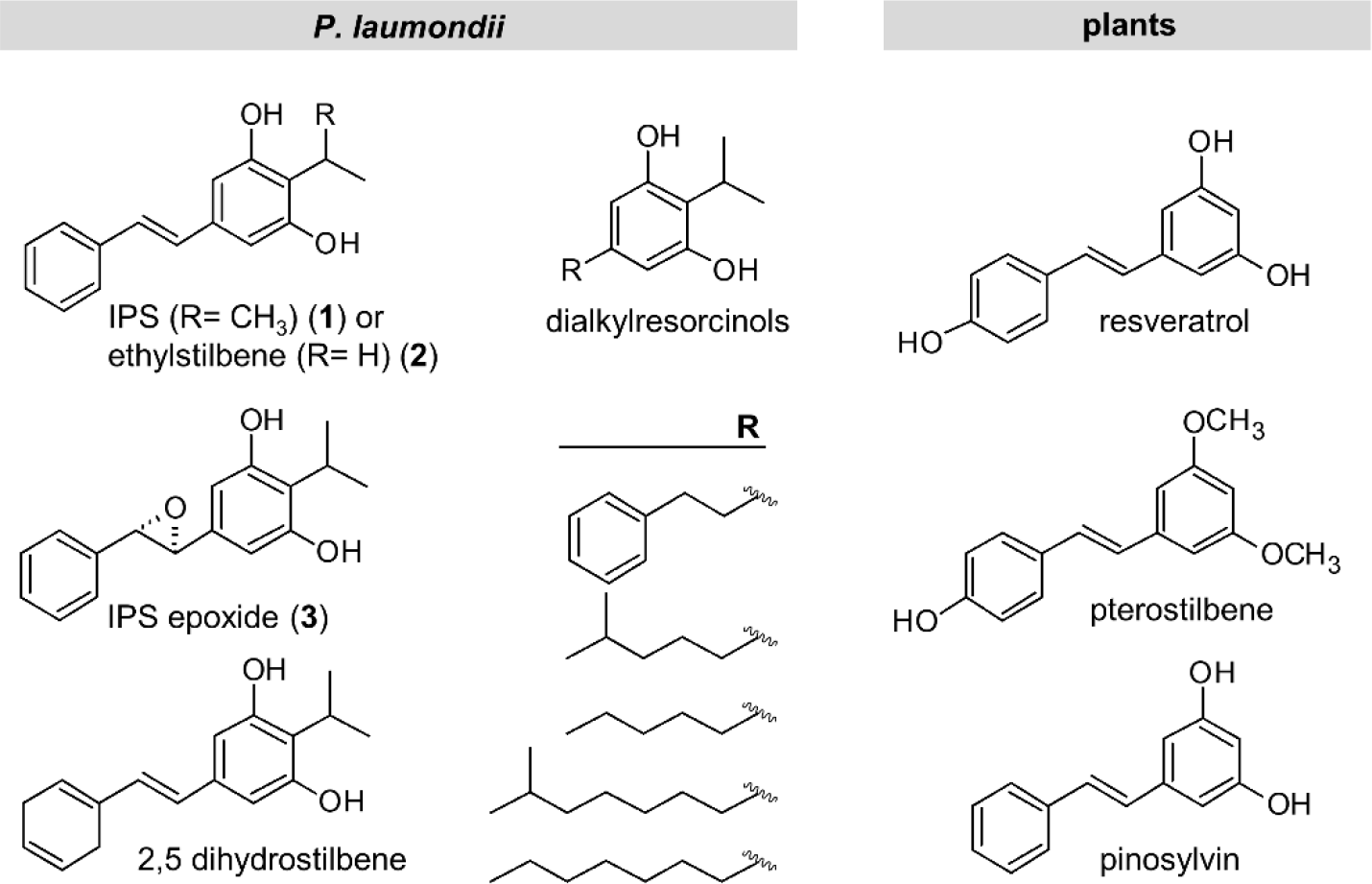
Selected stilbene and dialkylresorcinol (DAR) compounds produced by *P. laumondii* and stilbene representatives from plants.

The only non-plant stilbene is isopropylstilbene (IPS, Fig. 1, **1**) and derivatives thereof produced in *Photorhabdus* strains (Fig. 1).^5–11^ IPS is found in all *Photorhabdus* strains analyzed so far and can be regarded as a species-specific biomarker.^12^ *Photorhabdus* lives in mutualistic symbiosis with *Heterorhabditis* nematodes, acting together as an insect pathogen complex.^13–15^ It has been proven that IPS is required for nematode development, is important for the mutualistic symbiosis between bacterium and nematode, might act as live cycle signal for *Photorhabdus* itself and shows developmental and behavioral effects on other nematodes as well.^5,16,17^ Due to its antimicrobial activity, IPS might furthermore serve as a protection against microbial competitors during insect infection. Additionally, IPS inhibits protozoa and even the mammalian soluble epoxide hydrolase, which is involved in the arachidonic acid cascade and thus in several inflammatory processes.^5^ IPS is also known as tapinarof or benvitimod selectively modulating the cytokine cascade deep under the skin when applied as atopical cream.^18^ Within the last years it was developed as treatment against psoriasis, recently approved by the FDA and marketed as Vtama^®^.^5,18–20^

Interestingly, *Photorhabdus laumondii* utilizes a non-plant-like mechanism to produce IPS that relies on the condensation of two intermediates derived from different biosynthetic routes (Fig. 2b) which is crucial for obtaining the iso-branch.^5,6,21^ The corresponding genes are distributed within the genome and are encoded as stand-alone genes or short operons (*stlA* and *stlB, stlCDE* and *bkdABC*) (Fig. 2a). IPS-biosynthesis starts with the generation of cinnamic acid from phenylalanine catalyzed by a phenylalanine ammonia lyase (PAL, StlA). StlB is proposed to act as a CoA-ligase, activating cinnamic acid to cinnamoyl-CoA. The acyl moiety of cinnamoyl-CoA is suggested to be transferred on an acyl-carrier protein (ACP) followed by elongation, reduction and dehydration steps to obtain 5-phenyl-2,4-pentadienoyl-ACP. Since there are no ketosynthase, ketoreductase or dehydratase encoded in the *Photorhabdus* genomes in proximity with the already known genes, these reactions might be performed by enzymes from the fatty acid biosynthesis. To date, there is no proof for this hypothesis and furthermore, no acyltransferase transferring the acyl-moiety of cinnamoyl-CoA to the ACP (StlE) was identified so far. The dependence of these reactions on StlE was shown from analysis of a *ngrA* deletion.^5^ NgrA is an Sfp-type PPTase that is required for the biosynthesis of most PKS and non-ribosomal peptide synthetase derived peptides in *P. laumondii* TT01.^22^ The Michael acceptor 5-phenyl-2,4-pentadienoyl-StlE is further cyclized and aromatized with a β-ketoacyl isovalerate moiety derived from the branched-chain-α-keto acid dehydrogenase (BKD) complex (BkdABC), leading to IPS formation with the characteristic iso-branched side chain of stilbenes produced from *P. laumondii* (Fig. 2b).^5,6^ While the exclusive cyclization reaction was studied in the past with an *in vitro*-approach of the condensation (StlD) and subsequent aromatization (StlC) and with X-ray crystallography of the cyclase/ketosynthase (CYC/KS) StlD, the formation of the native substrate 5-phenyl-2,4-pentadienoyl-StlE remained elusive.^21^

**Figure 2.**
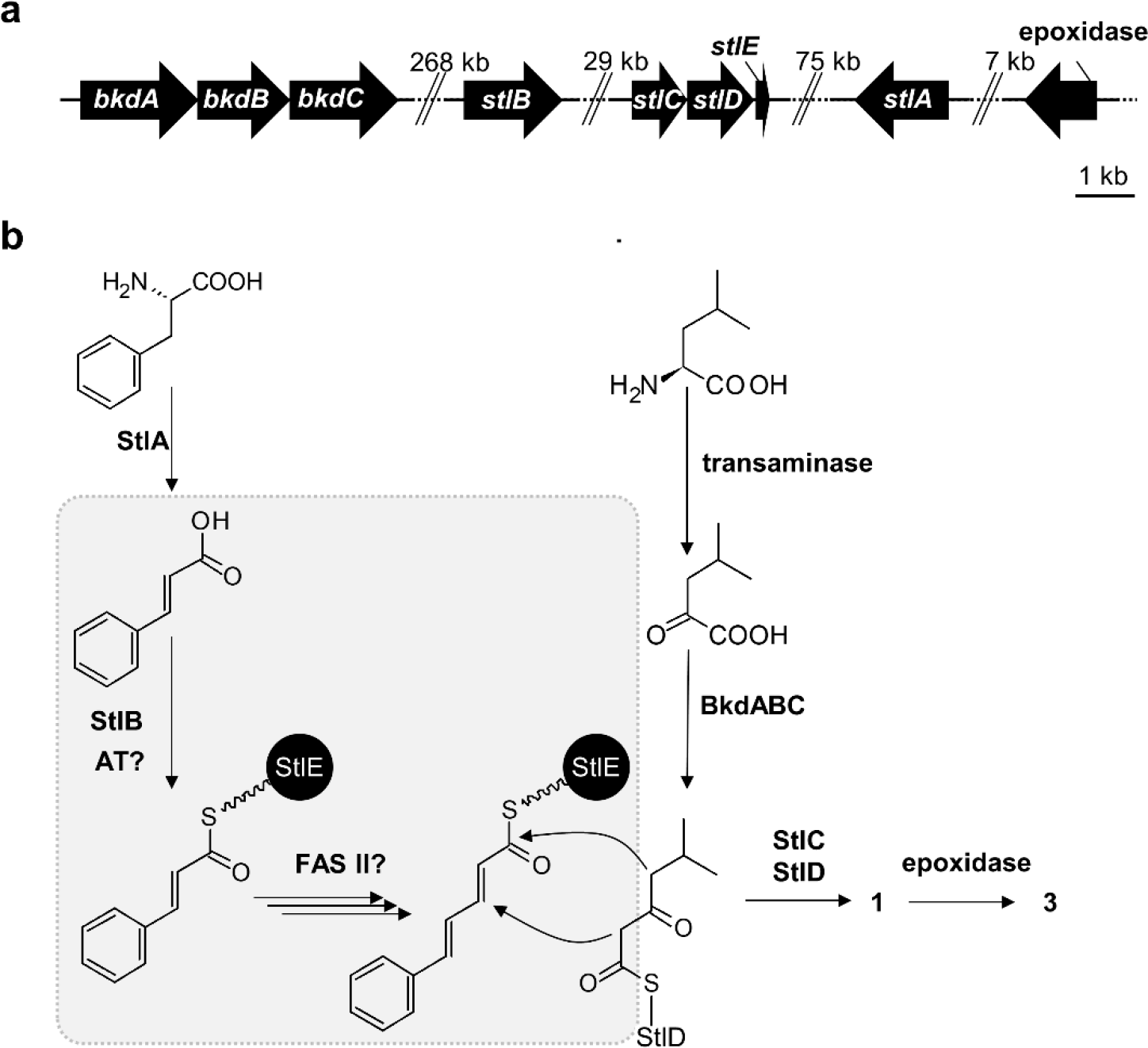
Genes involved in IPS production (**a**) and suggested IPS biosynthesis (**b**) in *P. laumondii*. BkdABC = branched-chain-α-keto acid dehydrogenase with ketosynthase (*plu1883-1885*), StlB = CoA ligase (*plu2134*), StlC = aromatase (*plu2163*), StlD = ketosynthase/cyclase (*plu2164*), StlE = acyl-carrier protein (*plu2165*), StlA = phenylalanine ammonium lyase (*plu2234*), epoxidase = *plu2236*Klicken oder tippen Sie hier, um Text einzugeben., AT = acyltransferase, FAS II = fatty acid synthase type II. Highlighted in grey are steps, which were investigated by *in vitro*-studies.

Here, we report the *in vitro* substrate characterization of StlB and furthermore the conversion of cinnamoyl-CoA to 5-phenyl-2,4-pentadienoyl-ACP by the orchestration of enzymes from the fatty acid biosynthesis pathway. Furthermore, we focused on IPS production in a heterologous *E. coli* host.

## 2. Results and Discussion

We started our investigations by characterizing StlB for its postulated function as a CoA ligase. Although cinnamic acid is the proposed natural substrate, cinnamoyl-CoA was hardly detectable during *in vitro* analysis with purified StlB via MALDI-MS when compared to CoA derivatives generated from fatty acid substrates including chain length from C_6_ to C_16_ (Fig. S1, Table S5).

To rule out any ionization problems of cinnamoyl-CoA compared to the fatty acid acyl-CoA thioesters, we analyzed reactions with cinnamic, caproic, heptanoic and palmitic acid with HPLC-UV-MS and were able to detect all formed CoA-esters by UV (260 nm, Fig. S2) and MS (Fig. S1) except cinnamoyl-CoA.

We further observed that StlB accepts some cinnamic acid derivatives but discriminates substituents in the *para*-position (Table S5). These findings are in accordance with previously conducted mutasynthesis experiments, where *para*-substituted cinnamic acid derivatives did not lead to novel IPS derivatives in *P. laumondii,* while meta-substituted substrates were incorporated by the producer.^23^

Since *stlB* is annotated as a FadD homolog, a fatty acyl-CoA synthetase, converting fatty acids to acyl-CoAs for further fatty acid degradation^24^, its influence on the fatty acid degradation profile was studied. Supplementation of TT01 cultures with azido-labelled palmitic acid (C_16_-fatty acid) and application of click-chemistry showed no difference between WT and *stlB* mutant since both showed similar degradation profiles in the HPLC-MS analysis(Fig. S3, S4). Thus, we could not verify the characteristics of StlB as a FadD enzyme.

The importance of *stlB* for IPS production was further assessed by introduction of an insertion mutation which resulted in the loss of the production (Fig. S5).

Since StlB produces less cinnamoyl-CoA than fatty acyl-CoA *in vitro*, we tested StlB for its activity as an adenylating enzyme, thus being capable of direct-loading of adenylated cinnamoyl-precursor to *holo*-StlE. Such enzymes are known as acyl-acyl carrier protein synthetases (AasS) or AMP-ligases.^25–28^ Hence, we purified the ACP StlE and transferred it into its *holo*-form with the PPTase Sfp.^29^ When we tested StlB for AasS activity in the presence of cinnamic acid and *holo*-StlE, we were indeed able to show the transferase reaction (Fig. 3a). Hence, we suggested StlB to substitute the “missing” acyltransferase responsible for cinnamic acid activation and StlE acylation to result in cinnamoyl-StlE. Furthermore, analysis of StlB with fatty acid substrates demonstrated acyl transfer of the fatty acid substrates and also acylation of AcpP from *E. coli* (Fig. S6). StlE and the *E. coli* derived AcpP share around 60 % sequence identity and around 80 % in the region of the helix II which is essential for the recognition and interaction with other proteins leading to similar interaction partners between those two ACPs.^30^

**Figure 3.**
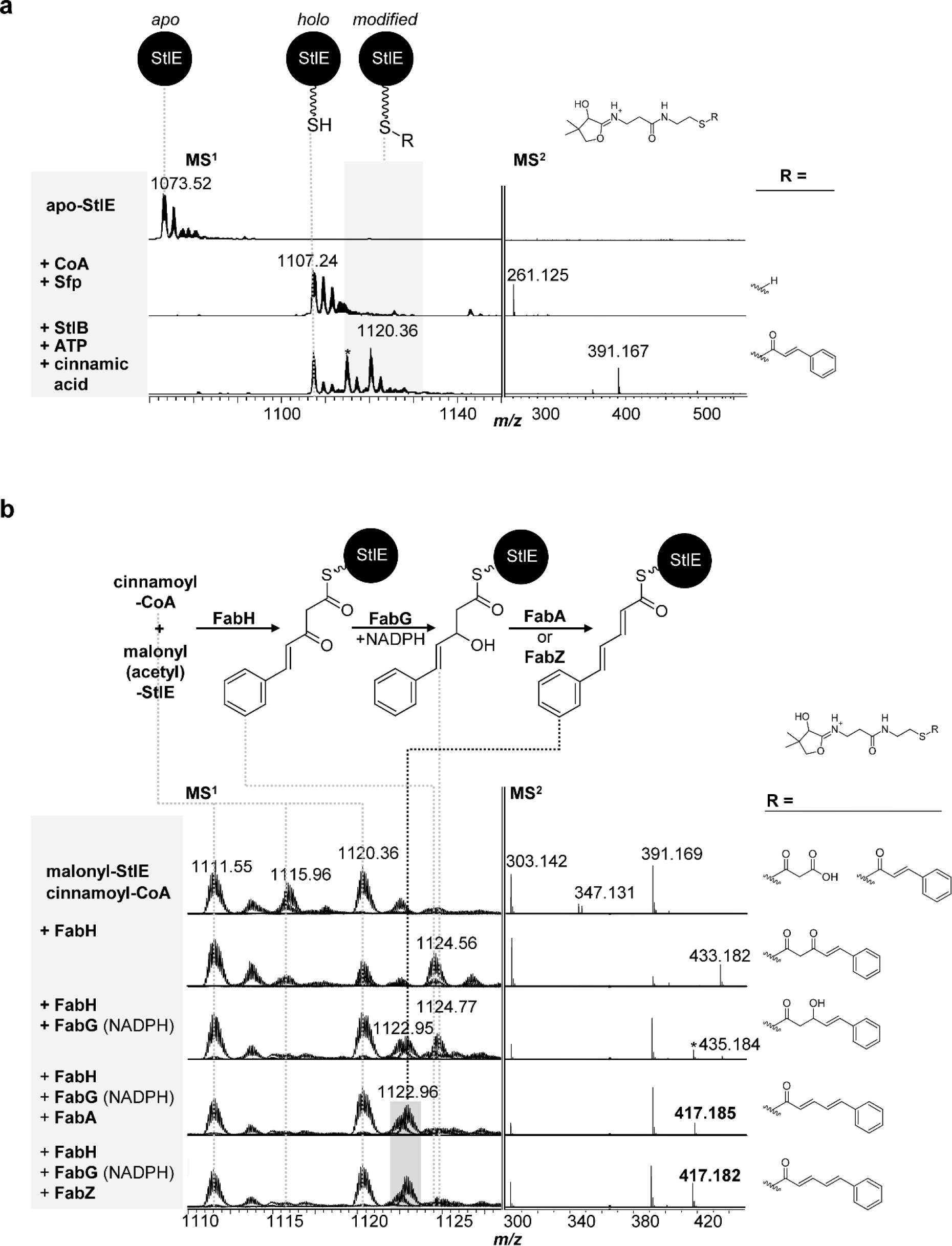
AasS loading of StlE catalyzed by StlB. *apo*-StlE was transferred to *holo* StlE with Sfp. *holo*-StlE was transferred to cinnamoyl-StlE by the AasS activity of StlB. Reaction marked with an asterisk indicates an unknown side product arising during overnight incubation (**a**). *In vitro* elongation, ketoreduction and dehydration reaction with FAS II enzymes FabH, FabG (+NADPH), FabA, FabZ from *P. laumondii* and cinnamoyl-CoA as starter and malonyl-ACP as elongation unit. Spectra where overlaid as described in the experimental section. Signal marked with an asterisk represents the dehydrated product without the addition of FabA/FabZ (**b**).

IPS biosynthesis requires the formation of 5-phenyl-2,4-pentadienoyl-StlE as substrate for the cyclization reaction catalyzed by the ketosynthase/cyclase StlD (see Fig. 2b).^5,6,21^ StlD only accepts unsaturated intermediates.^6,21^ Hence, the elongation of cinnamoyl-StlE and its subsequent reduction and dehydration is crucial for cyclization. Since none of the required enzymes, namely a ketosynthase, ketoreductase and a dehydratase, are encoded next to the known IPS biosynthesis enzymes, we tested enzymes from the fatty acid biosynthesis of *P. laumondii* as possible candidates for these reactions. For elongation reaction, we selected FabH, FabB and FabF as plausible ketosynthase candidates, as the special ketosynthase StlD is not able to fulfill this task. FabH is usually responsible for the initial elongation of acetyl-CoA starter with malonate as elongation units in FA biosynthesis, while FabB and FabF are responsible for elongation of longer acyl-ACP supported by distinct protein-protein interactions.^31,32^ In our *in vitro* analysis with purified enzymes (Fig. S7), only FabH was able to elongate the cinnamoyl-moiety with malonyl-ACP to its β-ketoacyl product, while FabB or FabF were not (Fig. S8). These findings contradict with the usual role of FabH as a priming ketosynthases involved in the initiation of the biosynthesis with a short chain acyl-CoA in the fatty acid metabolism. It is also in contrast to the ability of StlB to act as an AasS enzyme within the pathway that would not require an extra FabH for the procession of a CoA-bound moiety. However, we showed that in the *in vitro* approach cinnamoyl-CoA was sufficient for the procession into 5-phenyl-2,4-pentadienoyl-StlE since the cinnamoyl-moiety is loaded onto StlE (Fig. 3b, *m/z* 1120.36 [M+H]^+^) in that reaction setup and that further ketoreduction was performed by FabG and dehydration by FabA or FabZ (Fig. 3b).

Although the dehydrated product was formed even without the addition of either FabA nor FabZ, the amount of dehydrated product was increased upon the addition of a dehydratase (Fig. 3b). Furthermore, in the absence of the dehydratases, the amount of the dehydrated side product remained stable upon 2 h incubation time, which is equivalent to the incubation time used for the assay (Fig. S9).

With the gained knowledge from our *in vitro*-assays, we focused on the production of stilbene compounds in a heterologous host. So far, only dialkylresorcinol (DAR) compounds, lacking the cinnamoyl side chain, were detected in heterologous expression experiments with StlCDE-homologs.^33^ Having now identified all key players for stilbene biosynthesis, for the first time, we successfully produced IPS (**1**) in 0.61 ± 0.09 mg/L in a heterologous *E. coli* host (Fig. 4). The quantification was achieved by preparation of a standard curve of purified IPS from *P. laumondii* via HPLC-MS and cultivation of triplicates from the *E. coli* production strain. The production of **1** was achieved by co-expression of *stlCDE*, *stlB*, *plFabH*, *plfabG* and *plBkdABC* and cinnamic acid supplementation. It was not necessary to express *ngrA* since the *E. coli* PPTases were able to catalzye the modification of StlE from *apo-* to *holo-*form.

**Figure 4.**
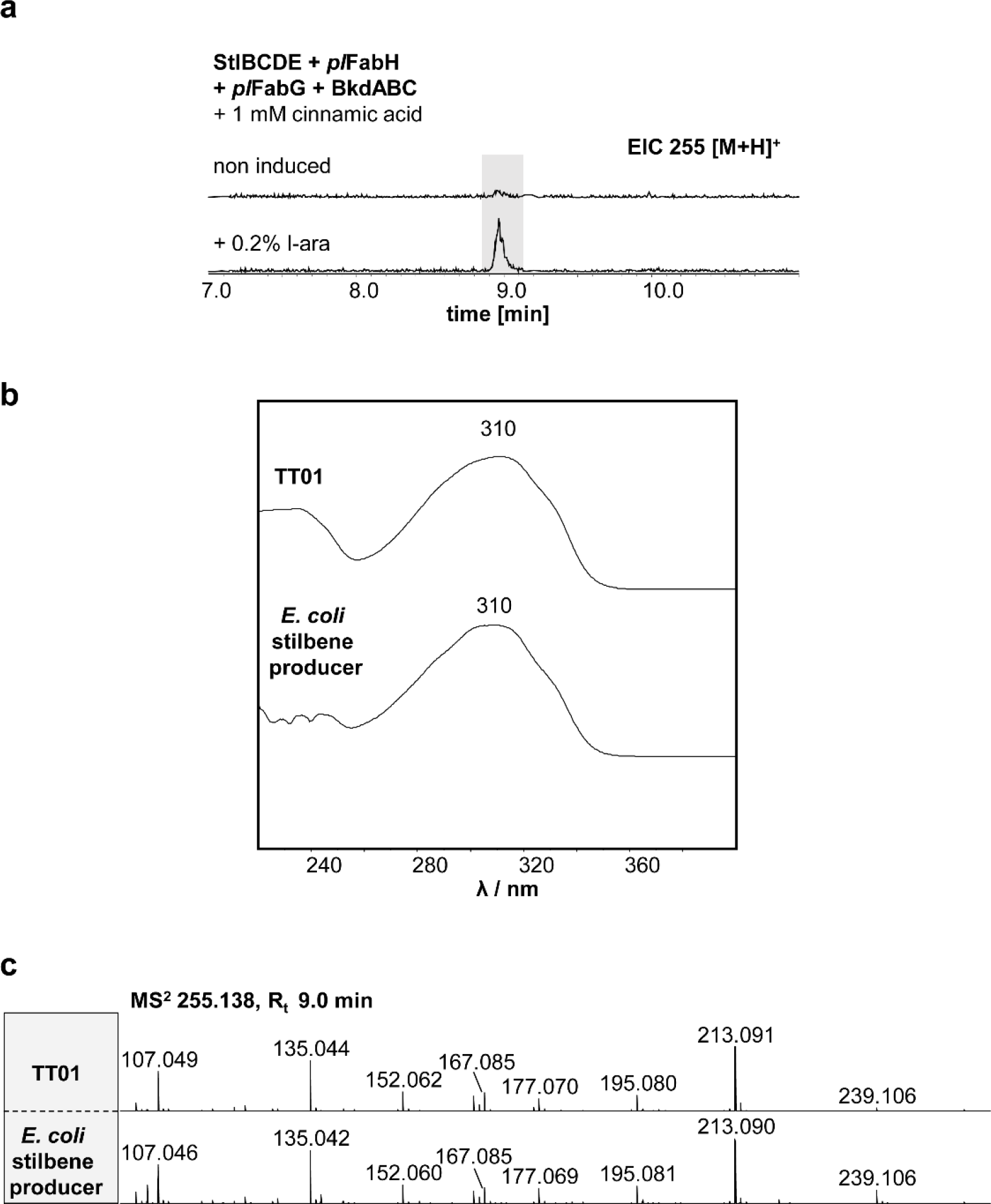
Heterologous stilbene production in *E. coli*. Analysis of IPS production after cultivation of *E. coli* DH10B with introduced *stlBCDE*, *pl*fabHbkdABC and *pl*FabG. The cultures were supplemented with 1 mM cinnamic acid. The crude extracts of the corresponding strains were analyzed by HPLC-UV-MS in positive mode. The EIC of IPS ([M+H]^+^ 255.138±0.005 Da) is depicted (a) and compared to the UV and MS^2^ data from IPS isolated from *P. laumondii* TT01 (b and c).

## 3. Conclusion

We revealed the missing parts of the IPS biosynthesis from *P. laumondii* to generate 5-phenyl-2,4-pentadienoyl-StlE, the required precursor for cyclization reaction by the KS/CYC StlD (Fig. 5). We demonstrated that StlB is able to acylate the cognate ACP StlE to form cinnamoyl-StlE and cinnamoyl-CoA. Cinnamoyl-CoA is further used by enzymes from FAS II biosynthesis for elongation, reduction and dehydration reaction, mediated by FabH, FabG and FabA or FabZ, respectively. While especially the role of FabH in the biosynthesis requires further investigations including structural biology data, our data shows that the IPS biosynthesis is another example of cross-talk between FAS II enzymes from primary metabolism and proteins from specialized metabolism. Similar cross-talk between specialized and primary metabolism and metabolites was described in the formation of the rhabdoplanins formed via an ‘Ugi-like’ reaction, including building blocks from primary metabolism and the specialized metabolite rhabduscin.^34^ Our results helped us to further select the minimal gene set needed for the production of isopropylstilbenes in a heterologous host. These efforts might be the starting point for further engineering strategies in order to produce additional stilbene derivatives in the future.

**Figure 5.**
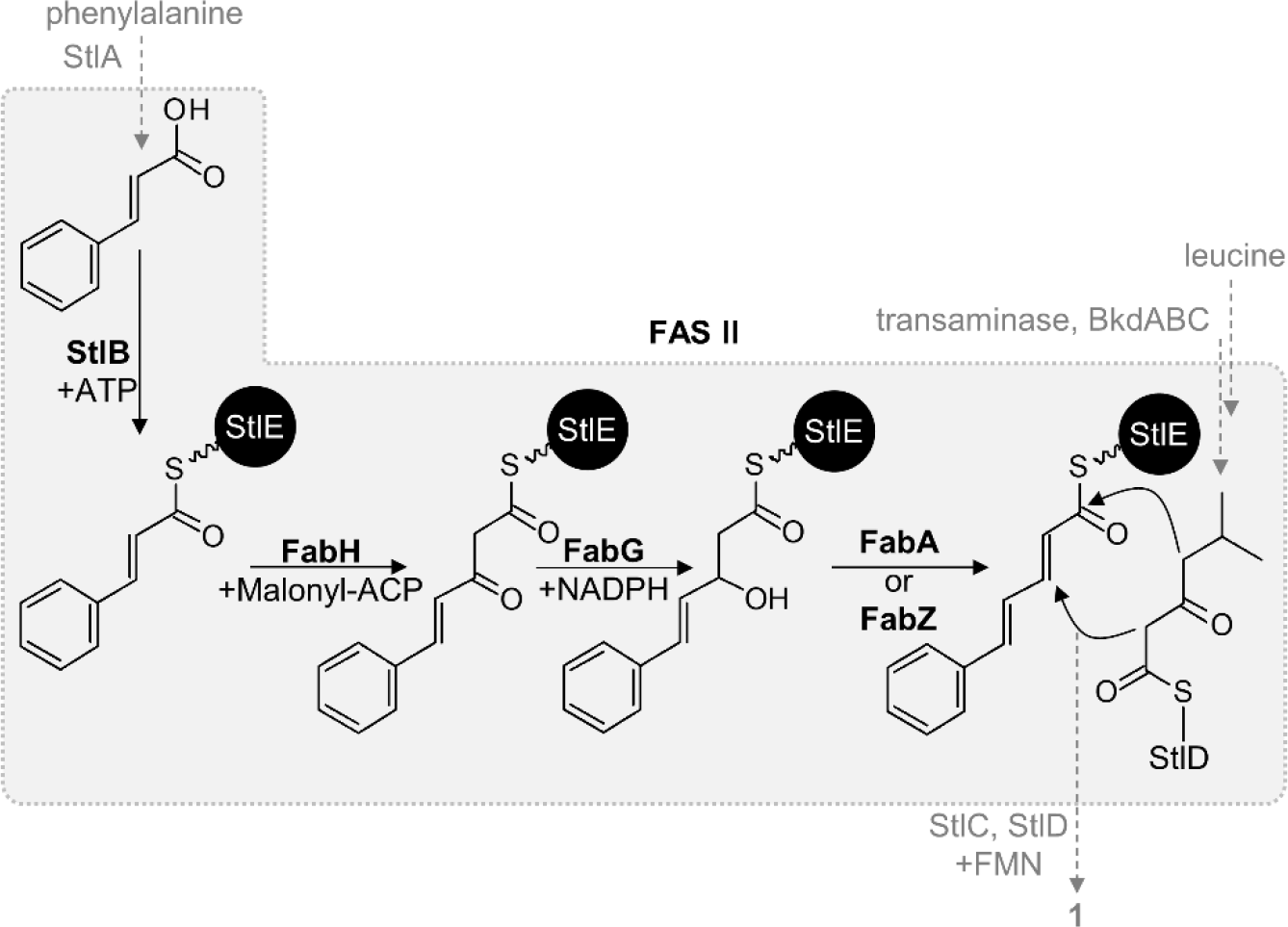
Extended IPS-biosynthesis. The cinnamoyl moiety of the adenylated precursor is loaded to StlE by the acyl-acyl-carrier protein synthetase (AasS) activity of StlB before being elongated with one malonate unit by FabH. The resulting β-ketoacyl product is further reduced and dehydrated by FabG and FabZ/FabA. StlB = AasS, FabH = ketosynthase; FabG = ketoreductase; FabA/FabZ = dehydratase; FAS II = fatty acid synthase type II, StlC = aromatase, StlD = ketosynthase/cyclase; BkdABC = branched-chain-α-keto acid dehydrogenase with ketosynthase (for further details see Fig. 2).

## 4. Experimental Section

### 4.1 Cloning

For isolation of genomic DNA from *P. laumondii,* Gentra Puregene Yeast/Bact Kit (Qiagen) was used. Polymerase chain reaction (PCR) was performed with Phire Hot Start II DNA polymerase (Thermo Scientific), Phusion High-Fidelity DNA Polymerase, or Q5 polymerase (New England Biolabs) according to manufacturers’ instructions. Oligonucleotides were purchased from Eurofins Genomics. The Invisorb Spin DNA Extraction Kit (Stratec) was used for DNA purification from agarose gels. Plasmid isolation was performed with the Invisorb Spin Plasmid Mini Two Kit (Stratec). Plasmid-backbone PCRs were restriction digested with *Dpn*I (New England Biolabs) following the manufacturers’ protocol. All used primers and templates and final plasmids are listed in Table S2 and S3. If not stated otherwise, all plasmids were cloned via Hot Fusion^35^ reaction with corresponding primers listed in Table S2. *plu2134* gene region was amplified using TS120 and TS121, generating a PCR product of 649 bp, which was digested (*Sac*I and *Sph*I) and ligated (T4 DNA ligase, Sigma) into the linearized vector (*Sac*I and *Sph*I) pDS132. *E. coli* S17λpir (pDS132_plu2134) and *E. coli* DH10B (remaining plasmids) were used as cloning strains and electroporated with desalted ligation/Hot Fusion reaction (MF-Millipore membrane, VSWP, 0.025 μm).

### 4.2 Generation of TT01 mutant

The *plu2134* (*stlB*)-insertion mutant of *P. laumondii* TT01 was created by conjugation with the *E. coli* S17λpir strain harboring pDS132_plu2134, which carries the first 578 bp of *plu2134*, as described earlier for creating insertion-mutants.^36^ As a result, in a homologous recombination event, the plasmid backbone is inserted within the gene of interest in the case of *plu2134*, resulting in a non-functional gene (*plu2134*-insertion, *P. laumondii* TT01::pDS132_*plu2134*).

Therefore, both strains, *P. laumondii* TT01 and *E. coli* S17λpir with pDS132_plu2134, were grown in 10 mL LB-medium (10 g/L tryptone, 5 g/L yeast extract and 5 g/L NaCl at pH 7.5) (*E. coli* S17λpir was supplemented with 35 μg/mL chloramphenicol) to an OD_600_ of 0.6-0.8. Cells of 1 mL culture were harvested, washed and resuspended in 400 μL LB and mixed in a ratio of 3:1 (75 μL TT01:25 μL *E. coli*) prior to pipetting them in one drop on an LB agar plate. After 1 d at 30°C, cells were resuspended in 1 mL LB. Insertion mutants were selected by streaking several dilutions of cell suspension on LB-Cm^R^-Rif^R^ (35 μg/mL and 100 μg/mL, respectively) agar plates. Positive insertion mutants were PCR-verified with primer TS122 and pDS132_rev leading to an 1147 bp product.

### 4.3 Click reaction and XAD-extraction of TT01 wildtype and mutants

The fatty acid degradation profile of *P. laumondii* was analyzed via feeding of azido palmitic acid followed by a click reaction with bicyclononyne (BCN).^37^ Therefore, pre-cultures of TT01 and mutants were grown overnight in LB-medium. The following day appropriate cell material was used for inoculation of the main cultures to an OD_600_ of 0.1, supplemented with 0.1 mM azido palmitic acid and incubated for 24 h at 30°C and 200 rpm. After 2 h, 4 h, 6 h, 10 h, 20 h and 24 h, 1 mL cell culture of each strain was harvested and stored at −20°C. The click reaction of (the degradated) azido palmitic acid was performed as previously described.^37^ Briefly, the cell pellet was resuspended in 500 μL 1 M NaOH and incubated for 1-2 h at 85°C and 750 rpm on a thermocycler. The solution was acidified with 100 μL 6 M HCl prior to extraction with 800 μl hexane. 600 μL of the organic layer was evaporated under reduced pressure. The click reaction was performed with 150 μL of 0.2 mM BCN (in ACN) and incubated overnight at RT. A 1:10 dilution of each click reaction was analyzed via HR-HPLC-MS. The expected masses of the click products are listed in Figure S3.

To obtain a comparable growth rate of wildtype and mutant strains, the cultivation was carried out without the addition of antibiotics. After 72 h, the plasmid insertion of *P. laumondii* TT01::P_DS132_*plu2134* was proven (no production of **1**) by HPLC-MS (Fig. S5). Therefore, the strain was cultivated in 10 mL LB as described above, with further supplementation of 2% Amberlite® XAD-16 (Sigma-Aldrich) beads. After 48 h, the XAD-beads were extracted with 10 mL MeOH, filtrated and dried under reduced pressure. Samples were further analyzed by HPLC-UV/MS.

### 4.4 Heterologous stilbene production

DH10B was transformed with the production plasmids listed in Table S4, respectively. For detection of **IPS**, extracts from 10 mL LB-culture of each strain were prepared upon XAD-bead addition. Therefore, the corresponding overnight pre-culture was used for inoculation of 10 mL LB-medium to an OD_600_ of 0.2 and supplemented with 0.2% l-arabinose, with and without 1 mM cinnamic acid (Sigma-Aldrich) (from 100 mM cinnamic acid stock solution in ethanol) and 2% Amberlite® XAD-16 (Sigma-Aldrich) resin. After 48 h, XAD-beads of each culture were extracted with 10 mL MeOH, filtered and dried under reduced pressure. For HR-HPLC-MS measurement, the samples were dissolved in 1 mL MeOH.

### 4.5 Protein production and purification

*E. coli* BL21 (DE3) was transformed with protein production plasmids (Table S3). Protein production was achieved using an auto-induction protocol.^5^ Therefore 1 L LB-medium was supplemented with 20 mL 50 × 5052 (25% glycerol, 2.5% glucose, and 10% α-lactose monohydrate), 50 mL 20 × NPS (1 M Na_2_HPO_4_, 1 M KH_2_PO_4_, and 0.5 M (NH_4_)_2_SO_4_), 1 mL 1 M MgSO_4_ *7 H_2_O and the respective antibiotics (35 μg/mL chloramphenicol, 50 μg/mL kanamycin, 50 μg/mL Spectinomycin, 20 μg/mL Gentamicin). This auto-induction medium was inoculated with the appropriate pre-culture and incubated at 37°C and 180 rpm to an OD_600_ = 0.4-0.8, cooled down for 15-45 min at 4°C and incubated overnight at 25°C and 180 rpm.

*Purification of Strep-tagged StlB, StlE, FabB, plFabF,pl FabH, plFabG, plFabA, plFabZ.* Cells were harvested and resuspended in Strep-tag binding buffer (400 mM NaCl, 50 mM Tris, pH 8, pH 8.5 StlB, pH 7.4 StlC) and with the addition of one tablet cOmplete Protease Inhibitor Cocktail (Roche) and lysozyme incubated for 20-50 min at 4°C. After cell lysis by sonication, cell debris was removed by centrifugation (35 min, 20,000 rpm, 4°C) and the supernatant was passed over a 5 mL StrepTrap HP column (GE Healthcare) by an ÄKTApurifier system (GE Healthcare). Proteins were eluted with Strep-tag elution buffer (400 mM NaCl, 50 mM Tris, 2.5 mM d-desthiobiotin, pH 8, pH 8.5 StlB, pH 7.4 StlC). All proteins were buffer exchanged in storage buffer (20 mM HEPES, 1 mM DTT, 25% (v/v) glycerol, pH 8), concentrated to 2-10 mg/ml using Amicon concentration devices (Merck, 3,000 or 10,000 Da cut-off filter) and stored in aliquots at −80 °C until needed. StlB was directly used after purification.

*Purification of His_6_-tagged* Sfp-SUMO. Cells were harvested and resuspended in His-tag binding buffer (300 mM NaCl, 20 mM Tris, 20 mM imidazol, pH 8.5) and with addition of one tablet cOmplete Protease Inhibitor Cocktail (Roche) and lysozyme incubated for 20-50 min at 4°C. After cell lysis by sonication, cell debris was removed by centrifugation (35 min, 20,000 rpm, 4°C) and the supernatant was passed over a 5 mL HisTrap FF column (GE Healthcare) by an ÄKTApurifier system (GE Healthcare). Proteins were eluted with His-tag elution buffer (300 mM NaCl, 20 mM Tris, 500 mM imidazol, pH 8.5) and buffer exchanged in storage buffer (20 mM HEPES, 1 mM DTT, 25% (v/v) glycerol, pH 8.5) using a PD-10 desalting column (Amersham Biosciences). Proteins were further concentrated to 2-10 mg/ml using Amicon concentration devices (Merck, 3,000 Da, 10,000 Da, or 30,000 Da cut-off filter, respectively) and stored in aliquots at −80 °C until needed.

Proteins were separated on 10-15% SDS-polyacrylamide gels and visualized with Coomassie Brilliant Blue G250.

### 4.6 *In vitro* reactions

#### 4.6.1 CoA ligase activity assay

Reaction was performed in reaction buffer (10 mM Tris, 1 mM DTT, pH 8.5) with 1 mM CoA, 2 mM ATP, 5 mM MgCl_2_, 1 mM acid-substrate (Table S5) (solubilized as 10 mM stock in ethanol), 25 μM StlB (in Strep-elution buffer) and incubated for 1 h at 37°C. Reactions were analyzed by HR-HPLC-MS.

#### 4.6.2 AasS activity of StlB

StlE (ACP) or *ec*ACP (AcpP) was buffer exchanged in reaction buffer (10 mM Tris, 1 mM DTT, pH 8.5) with PD SpinTrap G-25 columns (Amersham Biosciences) and reactions were performed using 50 μM ACP, 1 mM CoA, 0.5 μM Sfp-SUMO (in storage buffer), 2 mM ATP, 5 mM MgCl_2_, 1 mM acid-substrate (Table S5) (solubilized as 10 mM stock in ethanol), 5-15 μM StlB (in storage buffer) and incubated over night at 30°C. Reactions were analyzed by HR-HPLC-MS.

#### 4.6.3 Preparation of malonyl-StlE (elongation-ACP)

ACP StlE was buffer exchanged in reaction buffer (10 mM Tris, 1 mM DTT, pH 8.5) with PD SpinTrap G-25 columns (Amersham Biosciences). Reactions were performed using 50 μM StlE, 1 mM malonyl-CoA, 0.5 μM Sfp-SUMO (in storage buffer) and incubated for 1 h at 30°C. The reaction was analyzed by HR-HPLC-MS.

#### 4.6.4 Chain elongation, ketoreduction and dehydration with FAS II enzymes

Starter-CoA and elongation-ACP assay were mixed 1:2 and chain elongation reaction was performed by adding 10 μM FabH or FabB or FabF and further incubated for 1 h at 30°C, prior to adding 10 μM FabG and 1 mM NADPH for ketoreduction reaction and 10 μM FabA or FabZ for dehydration reaction.

All ACP-bound *in vitro* reactions were analyzed using HR-HPLC-MS. MS^1^ and MS^2^ BPC and TIC spectra were summarized according to the retention time of modified ACP. If the retention time of two ACP species differed more than 0.4 min, the separately summarized spectra were overlaid. Theoretical average masses of modified ACPs were predicted using IsotopePattern (Bruker).

### 4.7 HR-HPLC-UV/MS

Click-reactions and XAD extracts of TT01 and TT01 mutants as well as *in vitro*-assays of CoA ligase and ACP-bound reactions were analyzed via high resolution (HR)-HPLC-ESI-UV-MS using a Dionex Ultimate 3000 LC system (Thermo Fisher) equipped with a DAD (Impact II) or MWD (micrOTOF II)-3000 RS UV-detector (Thermo Fisher) and coupled to an Impact II or micrOTOF II electrospray ionization mass spectrometer (Bruker).

*Click-reactions and XAD extracts.* Separation of Click-reaction and XAD extracts was achieved on a C18 column (ACQUITY UPLC BEH, 50 mm × 2.1 mm × 1.7 μm, Waters) using H_2_O and ACN containing 0.1% (v/v) formic acid as mobile phases. HPLC was performed at a flow rate of 0.4 mL/min with 5% ACN equilibration (0-2 min) followed by a gradient from 5-95% ACN (2-14 min, 14-15 min 95% ACN) ending with a re-equilibration step of 5% ACN (15-16 min). For internal mass calibration, 10 mM sodium formate was injected. The HPLC-MS analysis was set to negative mode with a mass range of *m/z* 100-1200 with and an UV at 190-800 nm.

*CoA ligase-reaction* (Impact II). Analysis of CoA-esters was performed on a C18 column (ACQUITY UPLC BEH, 50 mm × 2.1 mm × 1.7 μm, Waters) using MeOH and 50 mM ammonium acetate as mobile phases. HPLC was performed at a flow rate of 0.3 mL/min with 5% ACN equilibration (0-2 min), followed by a gradient from 5-95% ACN (2-14 min, 14-15 min 95% ACN) ending with a re-equilibration step of 5% ACN (15-16 min). For internal mass calibration, 10 mM sodium formate was injected.

*ACP-bound reactions* (micrOTOF II). Analysis of ACP derivatives was performed with a C3 column (Zorbax 300SB-C3 300Å, 150 mm × 3.0 mm × 3.5 μm, Agilent). ACN and H_2_O containing 0.1% (v/v) formic acid were used as mobile phases at a flow rate of 0.6 mL/min. HPLC was performed with 30% ACN equilibration (0-1.5 min), followed by a gradient from 30-65% ACN (1.5-27 min) and a further elution step with 95% ACN (27-30 min). For internal mass calibration, an ESI-L Mix (Agilent) was injected.

For data analysis of UV-MS-chromatograms, Compass DataAnalysis 4.3 (Bruker) was used. The theoretical average masses of StlE and *ec*ACP were calculated using Compass IsotopePattern 3.0 (Bruker).

### 4.8 MALDI-MS

*In vitro* CoA ligase reactions were analyzed via MALDI-MS on an LTQ Orbitrap XL instrument (Thermo Fisher Scientific, Inc.) equipped with a nitrogen laser at 337 nm. The samples were mixed with the matrix (3 mg/mL in 75% ACN with 0.1% TFA) while spotting them onto a polished stainless steel MALDI target in a 1:3 ratio (0.5 μL sample/1.5 μL matrix) with addition of 0.25 μL ProteoMass Normal Mass Calibration Mix (1:10 dilution, ProteoMass™ MALDI Calibration Kit, Sigma-Aldrich) for acquisition of high resolution spectra. Mass spectra were acquired in positive ion mode (FTMS) over a range of 700-1200 *m/z*. MS^2^ experiments were performed in ITMS mode with ±0.1 Da mass accuracy window, wideband activation and 23 eV collision energy. Data analysis was performed using Xcalibur 2.0.7 (Thermo Fisher Scientific, Inc.) by averaging 30-100 consecutive laser shots.

## Acknowledgements

Work in the Bode laboratory was supported by the BMBF project RhabdoFerm (031B0856A) and in part by the ERC advanced grant SYNPEP (835108) and the LOEWE Schwerpunkt MegaSyn, funded by the State of Hesse. We acknowledge the Deutsche Forschungsgemeinschaft for funding of the Impact II qTof mass spectrometer (Grant INST 161/810-1). We thank Alexander Perez for providing the azido-C_16_ fatty acid, Alexander Rill for providing the plasmid backbones for the heterologous production, Carsten Kegler and Charles O. Rock for providing the plasmids for Sfp- and AcpP-production, respectively. We further thank Michael Karas for MALDI access.

