## Supplemental data for "Biosynthesis of the Multifunctional Isopropylstilbene in *Photorhabdus laumondii* Involves Cross-Talk between Specialized and Primary Metabolism"

### SUPPORTING INFORMATION

#### Supporting Tables

**Table S1.** Strains used in this study.

| Strain | Genotype | Reference |
|---|---|---|
| *E. coli* DH10B | $F^-$ *araDJ39* Δ(*ara*, *leu*)7697 Δ*lacX74* *galU* *galK* *rpsL* *deoR* *ϕ*8O*dlac*ZΔM15 *endA*I *nupG* *recAl* *mcrA* Δ(*mrr hsdRMS mcrBC*) | Invitrogen |
| *E. coli* BL21 (DE3) Star | $F^-$, *ompT*, *gal*, *dcm*, *hsdSB*($r_B$- $m_B$-), *lon*, *λ(DE3* [*lacI*, *lacUV5-T7*, *gene1*, *ind1*, *sam7*, *nin5*]) | Invitrogen |
| S17-1 λ pir | $Tp^r$ $Sm^r$ *recA thi hsdRM*$^+$, RP4-2-Tc::Mu-Km::Tn7, *λpir* phage lysogen | [1] |
| *P. laumondii* TT01 | wild type, $Rif^R$ (spontaneous) | [2] |
| *P. laumondii* TT01::pDS132_*plu2134* | wild type with a $P_{DS132}$ insertion in *plu2134*, $Cm^R$ | this work |

**Table S2.** Oligonucleotides used for plasmid construction and verification. Overhangs are underlined.

| Plasmid | Oligonucleotide 5` to 3` | | Template |
|---|---|---|---|
| pDS132_plu2134 | TS120 | AAAGTGAGCTCCTGGCTAAAACATTATCCGG | *P. laumondii* TT01 |
| | TS121 | TGCCACCAGTATATTGCAGA | |
| pACYC_plu2134_strep (*stlB*) | GG185 | TTCGAAAAAGAGAACCTATACTTCCAGGGAGAGAAAGTCTGGCTAAAACATTATC | *P. laumondii* TT01 |
| | GG186 | AGCGGTGGCAGCAGCCTAGGTTAACTATGCTACATTCCTGACCTTTTC | |
| | GG68 | TCCCTGGAAGTATAGGTTCTCTTTTTCGAACTGCGGGTGGCTCCACATGGTATATCTCCTTATTAAAGTTAAAC | pACYC Duet-1 |
| | SW1 | TTAACCTAGGCTGCTG | |
| pCOLA_plu2134_strep (*stlB*) | GG185 | TTCGAAAAAGAGAACCTATACTTCCAGGGAGAGAAAGTCTGGCTAAAACATTATC | *P. laumondii* TT01 |
| | GG186 | AGCGGTGGCAGCAGCCTAGGTTAACTATGCTACATTCCTGACCTTTTC | |
| | GG68 | TCCCTGGAAGTATAGGTTCTCTTTTTCGAACTGCGGGTGGCTCCACATGGTATATCTCCTTATTAAAGTTAAAC | pCOLA Duet-1 |
| | SW1 | TTAACCTAGGCTGCTG | |
| pACYC_fabH_Strep | SK18 | GAGAACCTATACTTCCAGGGATATACAAAAATTTTAGGTACAGGTAGTTATC | *P. laumondii* TT01 |
| | SK22 | GATTACTTTCTGTTCGACTTAAGCATTATTAAAAACGTATCAGTGCCG | |
| | GG68 | TCCCTGGAAGTATAGGTTCTCTTTTTCGAACTGCGGGTGGCTCCACATGGTATATCTCCTTATTAAAGTTAAAC | pACYC Duet-1 |
| | SW1 | TTAACCTAGGCTGCTG | |
| pCOLA_bkdABC_Strep | SK23 | GAGAACCTATACTTCCAGGGAATAGACGTACAGATCAATGAAGTACC | *P. laumondii* TT01 |
| | SK24 | GATTACTTTCTGTTCGACTTAAGCATTATTACTGTTTTGCAACCACG | |
| | GG68 | TCCCTGGAAGTATAGGTTCTCTTTTTCGAACTGCGGGTGGCTCCACATGGTATATCTCCTTATTAAAGTTAAAC | pCOLA Duet-1 |
| | SW1 | TTAACCTAGGCTGCTG | |
| pCDF_stlB_Strep | SK25 | GAGAACCTATACTTCCAGGGAGAGAAAGTCTGGCTAAAACATTATC | *P. laumondii* TT01 |
| | SK26 | GATTACTTTCTGTTCGACTTAAGCATTACTATGCTACATTCCTGACCTTTTC | |
| | GG68 | TCCCTGGAAGTATAGGTTCTCTTTTTCGAACTGCGGGTGGCTCCACATGGTATATCTCCTTATTAAAGTTAAAC | pCDF Duet-1 |
| | SW1 | TTAACCTAGGCTGCTG | |
| pCDF_stlB_Strep_ StlCDE | SK27 | GTTAAGTATAAGAAGGAGATATACATATGAAACGTGTTCTTGTAGTGTCG | *P. laumondii* TT01 |
| | SK28 | GTGGCAGCAGCCTAGGTTAATTAGTTAGCGGATTCCAACTTTG | |
| | GG108 | CATATGTATATCTCCTTCTTATACTTAAC | pCDF_stlB_Strep |
| | GG109 | TAATTAACCTAGGCTGCTGCCAC | |
| pACYC_fabH_Strep_ fabG | SK29 | GTTAAGTATAAGAAGGAGATATACATATGAGATTAGATGGAAAGATTGCATTAG | *P. laumondii* TT01 |
| | SK30 | GTGGCAGCAGCCTAGGTTAATTAGTTCATATACATGCCGCC | |
| | GG108 | CATATGTATATCTCCTTCTTATACTTAAC | pACYC_fabH_ Strep |
| | GG109 | TAATTAACCTAGGCTGCTGCCAC | |
| pACYC_darABC (*stlCDE*) | GG1 | TATAAGAAGGAGATATACATATGAAACGTGTTCTTGTAGTG | *P. laumondii* TT01 |
| | GG8 | GTTTCTTTACCAGACTCGAGTTAGTTAGCGGATTCCAAC | |
| | GG3 | CTCGAGTCTGGTAAAGAAAC | pACYC Duet-1 |
| | GG4 | CATATGTATATCTCCTTCTTATACTTAAC | |
| pCDF_fabH_Strep | SK18 | GAGAACCTATACTTCCAGGGATATACAAAAATTTTAGGTACAGGTAGTTATC | *P. laumondii* TT01 |
| | SK19 | AGCGGTGGCAGCAGCCTAGGTTAATTAAAAACGTATCAGTGCCG | |
| | GG68 | TCCCTGGAAGTATAGGTTCTCTTTTTCGAACTGCGGGTGGCTCCACATGGTATATCTCCTTATTAAAGTTAAAC | pCDF Duet-1 |
| | SW1 | TTAACCTAGGCTGCTG | |
| pCDF_fabB_Strep | SK1 | GAGAACCTATACTTCCAGGGAAGGTATTTAATGAAGCGCGTAG | *P. laumondii* TT01 |
| | SK2 | AGCGGTGGCAGCAGCCTAGGTTAATTAAGCTGAATACTTGCTCATCAC | |
| | GG68 | TCCCTGGAAGTATAGGTTCTCTTTTTCGAACTGCGGGTGGCTCCACATGGTATATCTCCTTATTAAAGTTAAAC | pCDF Duet-1 |
| | SW1 | TTAACCTAGGCTGCTG | |

| | | | |
|---|---|---|---|
| pCDF_fabF_Strep | SK3 | GAGAACCTATACTTCCAGGGATCTAAGCGTCGAGTAGTTG | *P. laumondii* TT01 |
| | Sk4 | AGCGGTGGCAGCAGCCTAGGTTAATTAATCAGGTTTAATCTTACGG | |
| | GG68 | TCCCTGGAAGTATAGGTTCTCTTTTTCGAACTGCGGGTGGCTCCACATGGTATATCTCCTTATTAAAGTTAAAC | pCDF Duet-1 |
| | SW1 | TTAACCTAGGCTGCTG | |
| pACYC_fabG_Strep | SK5 | GAGAACCTATACTTCCAGGGAAGATTAGATGGAAAGATTGCATTAG | *P. laumondii* TT01 |
| | SK6 | AGCGGTGGCAGCAGCCTAGGTTAATTAGTTCATATACATGCCGC | |
| | GG68 | TCCCTGGAAGTATAGGTTCTCTTTTTCGAACTGCGGGTGGCTCCACATGGTATATCTCCTTATTAAAGTTAAAC | pACYC Duet-1 |
| | SW1 | TTAACCTAGGCTGCTG | |
| pCOLA_fabA_Strep | SK7 | GAGAACCTATACTTCCAGGGAGTTGATAAACTTAAATCCTACACAAAAG | *P. laumondii* TT01 |
| | SK8 | AGCGGTGGCAGCAGCCTAGGTTAATTAGAAAGCGCTGGTATCTTTAAAC | |
| | GG68 | TCCCTGGAAGTATAGGTTCTCTTTTTCGAACTGCGGGTGGCTCCACATGGTATATCTCCTTATTAAAGTTAAAC | pCOLA Duet-1 |
| | SW1 | TTAACCTAGGCTGCTG | |
| pCOLA_fabZ_Strep | SK9 | GAGAACCTATACTTCCAGGGAAGTGATAATCATACTCTGCACATTG | *P. laumondii* TT01 |
| | SK10 | AGCGGTGGCAGCAGCCTAGGTTAACTAAACCTCACGACGGC | |
| | GG68 | TCCCTGGAAGTATAGGTTCTCTTTTTCGAACTGCGGGTGGCTCCACATGGTATATCTCCTTATTAAAGTTAAAC | pCOLA Duet-1 |
| | SW1 | TTAACCTAGGCTGCTG | |
| pSEVA221-ara | AR585 | GTCGTGACTGGGAAAACC | pSEVA221 + araC Pbad fragment |
| | AR586 | TCCTGTGTGAAATTGTTATCC | |
| pSEVA631-ara | AR585 | GTCGTGACTGGGAAAACC | pSEVA631 + araC Pbad fragment |
| | AR586 | TCCTGTGTGAAATTGTTATCC | |
| pSEVA221-fabH-bkdABC-fabG | SK188 | AGAAAGAGGGGAAATACTAGATGTATACAAAAATTTTAGGTACAG | pSEVA221-ara |
| | SK189 | TTAATGCTTGTTAAAAACGTATCAGTGCC | |
| | SK190 | ACGTTTTTAACAAGCATTAAGCGCAATAAC | |
| | SK192 | AAGCTGTGTCTTACTGTTTTGCAACCAC | |
| | SK193 | AAAACAGTAAGACACAGCTTCACTAATAAC | |
| | SK194 | AGGGTTTTCCCAGTCACGACTTAGTTCATATACATGCCG | |
| | AR585 | GTCGTGACTGGGAAAACC | |
| | AR587 | CTAGTATTTCCCCTCTTTCTC | |
| pSEVA631-stlBCDE | SK184 | AGAAAGAGGGGAAATACTAGTTGGAGAAAGTCTGGCTAAAAC | pSEVA631-ara |
| | SK185 | ATCGGCACCTCTATGCTACATTCCTGACC | |
| | SK186 | TGTAGCATAGAGGTGCCGATGAATGGCTC | |
| | SK187 | AGGGTTTTCCCAGTCACGACTTAGTTAGCGGATTCCAACTTTGAAC | |
| | AR585 | GTCGTGACTGGGAAAACC | |
| | AR587 | CTAGTATTTCCCCTCTTTCTC | |
| araC-Pbad | | AAGCGGATAACAATTTCACACAGGATCATGACAACTTGA<br>CGGCTACATCATTCACTTTTTCTTCCAAACCGGCACGGA<br>ACTCGCTCGGGCTGGCCCCGGTGCATTTTTTAAATACCC<br>GCGAGAAATAGAGTTGATCGTCAAAACCAACATTGCGAC<br>CGACGGTGGCGATAGGCATCCGGGTGGTGCTCAAAAGC<br>AGCTTCGCCTGGCTGATACGTTGGTCCTCGCGCCAGCT<br>TAAGACGCTAATCCCTAACTGCTGGCGGAAAAGATGTGA<br>CAGACGCGACGGCGACAAGCAAACATGCTGTGCGACGC<br>TGGCGATATCAAAATTGCTGTCTGCCAGGTGATCGCTGA<br>TGTACTGACAAGCCTCGCGTACCCGATTATCCATCGGTG<br>GATGGAGCGATCCGTTAATCGCTAGCATGCGCCGCAGT<br>AACAATTGCTCAAGCAGATTTATCGCCAGCAGCTCCGAA<br>TAGCGCCCTTCCCCTTGCCCGGCGTTAATGATTTGCCCA<br>AAAAAGTCGCTGAAATGCGGCTGGTGCGCTTCATCCGG<br>GCGAAAGAACCCCGTATTGGCAAATATTGACGGCCAGTT<br>AAGCCATTCATGCCAGTAGGCGCGCGGACGAAAGTAAA<br>CCCACTGGTGATACCATTCGCGAGCCTCCGGATGACGA<br>CCGTAGTGATGAATCTCTCCTGGCGGGAACAGCAAAATA<br>TCACCCGGTCGGCAAACAAATTCTCGTCCCTGATTTTTC<br>ACCACCCCCTGACCGCGAATGGTGAGATTGAGAATATAA<br>CCTTTCATTCCCAGCGGTCGGTCGATAAAAAAATCGAGA<br>TAACCGTTGGCCTCAATCGGCGTTAAACCCGCCACCAGA<br>TGGGCATTAAACGAGTATCCCGGCAGCAGGGGATCATTT<br>TGCGCTTCAGCCATATTCACCACCCTGAATTGACTCTCTT | Purchased from Twist Bioscience |

| | | |
|---|---|---|
| | | CCGGGCGCTATCATGCCATACCGCGAAAGGTTTTGCAC CATTCGATGGCGCGCCGCCATTGGACCAAAACGAAAAA AGGCCCCCCTTTCGGGAGGCCTCTTTTCTGGAATTTGGT ACCGAGGCTTAACGATCGTTGGCTGAGAAACCAATTGTC CATATTGCATCAGACATTGCCGTCACTGCGTCTTTTACTG GCTCTTCTCGCTAACCAAACCGGTAACCCCGCTTATTAA AAGCATTCTGTAACAAAGCGGGACCAAAGCCATGACAAA AACGCGTAACAAAAGTGTCTATAATCACGGCAGAAAAGT CCACATTGATTATTTGCACGGCGTCACACTTTGCTATGC CATAGCATTTTTATCCATAAGATTAGCGGATCCTACCTGA CGCTTTTTATCGCAACTCTCTACTGTTTCTCCATACCCGA GCTGTCACCGGATGTGCTTTCCGGTCTGATGAGTCCGT GAGGACGAAACAGCCTCTACAAATAATTTTGTTTAATACT AGAGAAAGAGGGGAAATACTAG |
| Verification of *P. laumondii* TT01 *plu2134*- insertion mutant | | |
| **Oligonucleotide 5` to 3`** | | |
| TS121 | TCTCAGCTAGCAGCATCAAT | |
| pDS132_rev | GATCGATCCTCTAGAGTCGACCT | |

**Table S3.** Plasmids used in this study.

| Plasmid | Genotype | Reference |
|---|---|---|
| pACYC Duet-1 | P15A ori, T7*lac* promoter, $Cm^r$ | Novagen |
| pCDF Duet-1 | CDF ori, T7*lac* promoter, $Sm^r$ | Novagen |
| pCOLA Duet-1 | ColA ori, T7*lac* promoter, $Km^r$ | Novagen |
| pCATI4 | ColA ori, T7*lac* promoter, *cherry*, $Km^r$ | [3] |
| pDS132 | R6Kγ ori, *oriT*, *sacB, traJ*, *mob RP4*, $Cm^r$ | [4] |
| pSUMO_sfp | ColE1 ori, T7*lac* promoter, *sumo*, *sfp*, $Km^r$ | [5] |
| pET15b-AcpP | pBR322 ori, T7*lac* promoter, *acpP*, $Amp^r$ | Charles. O. Rock |
| pDS132_plu2134 | Insertion plasmid based on pDS132 with 578 bp of *plu2134*, $Cm^r$ | this work |
| pACYC_plu2134_strep | P15A ori, T7*lac* promoter, *plu2134 (stlB)*, $Cm^r$ | this work |
| pCOLA_plu2134_strep | ColA ori, T7*lac* promoter, *plu2134 (stlB)*, $Km^r$ | this work |
| pCOLA_darC_strep | ColA ori, T7*lac* promoter, *plu2165* (*stlE*), $Km^r$ | this work |
| pCDF_darA_Strep_BC (*stlCDE*) | CDF ori, T7*lac* promoter, *plu2163-2165* (*stlCDE*) $Sm^r$ | this work |
| pACYC_darA_Strep_B (*stlCD*) | P15A ori, T7*lac* promoter, *plu2163, plu2164* (*stlCD*), $Cm^r$ | this work |
| pACYC_darA_Strep (*stlC*) | P15A ori, T7*lac* promoter, *plu2163* (*stlC*), $Cm^r$ | this work |
| pCDF_darC_Strep (*stlE*) | CDF ori, T7*lac* promoter, *plu2165* (*stlE*), $Sm^r$ | this work |
| pCDF_darC_Strep_darB (*stlDE*) | CDF ori, T7*lac* promoter, *plu2164* (*stlD*) *plu2165* (*stlE*), $Sm^r$ | this work |
| pACYC_darABC (*stlCDE*) | P15A ori, T7*lac* promoter, *plu2163-2165* (*stlC*), $Cm^r$ | this work |
| pCDF_fabH_Strep | CDF ori, T7*lac* promoter, *plu2835*, $Sm^r$ | this work |
| pCDF_fabB_Strep | CDF ori, T7*lac* promoter, *plu3184*, $Sm^r$ | this work |
| pCDF_fabF_Strep | CDF ori, T7*lac* promoter, *plu2831*, $Sm^r$ | this work |
| pACYC_fabG_Strep | P15A ori, T7*lac* promoter, *plu2833*, $Cm^r$ | this work |
| pCOLA_fabA_Strep | ColA ori, T7*lac* promoter, *plu1772*, $Km^r$ | this work |
| pCOLA_fabZ_Strep | ColA ori, T7*lac* promoter, *plu0683*, $Km^r$ | this work |
| pACYC_fabH_Strep | P15A ori, T7*lac* promoter, *plu2835*, $Cm^r$ | this work |
| pCOLA_bkdABC_Strep | ColA ori, T7*lac* promoter, *plu1883-1885*, $Km^r$ | this work |
| pSEVA221 | RK2 ori, $Km^r$ | [6] |
| pSEVA631 | pBBR1 ori, $Gm^r$ | [6] |
| pSEVA221_ara | RK2 ori, ara promoter, $Km^r$ | Alexander Rill |
| pSEVA631_ara | pBBR1 ori, ara promoter, $Gm^r$ | Alexander Rill |
| pSEVA221-fabH-bkdABC-fabG | RK2 ori, ara promoter, *plu2835* (*fabH*), *plu1883- 1885* (*bkdABC*), *plu2833* (*fabG*), $Km^r$ | this work |
| pSEVA631-stlBCDE | pBBR1 ori, ara promoter, *plu2134 (stlB)*, *plu2163-2165* (*stlCDE*), $Gm^r$ | this work |

**Table S4.** *E. coli* DH10B strain used for heterologous stilbene production.

| Name | Introduced plasmids |
|---|---|
| StlBCDE+*pl*FabH+BkdABC+*pl*FabG | pSEVA221-fabH-bkdABC-fabG<br>pSEVA631-stlBCDE |

**Table S5.** Substrate specificity of StlB from *P. laumondii* TT01 by its CoA ligase reaction with substrates **1-14.** CoA ester formation was analyzed via MALDI-MS (Fig. S1).

| substrate | Compound name | structure | accepted |
|---|---|---|---|
| **1** | Cinnamic acid | 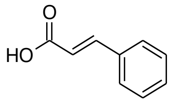 | + |
| **2** | Phenylpropanoic acid | 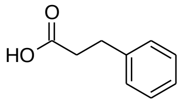 | + |
| **3** | 3-Chlorocinnamic acid | 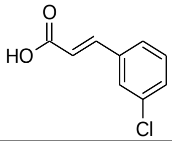 | + |
| **4** | 4-Chlorocinnamic acid | 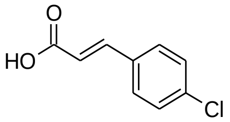 | - |
| **5** | Coumaric acid | 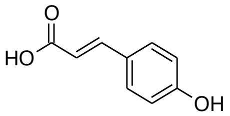 | - |
| **6** | Benzoic acid | 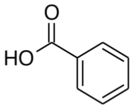 | - |
| **7** | Propionic acid | 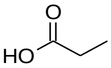 | - |
| **8** | Acetic acid | 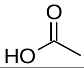 | - |
| **9** | Aminomalonic acid | 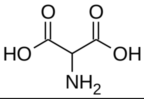 | - |
| **10** | Hexa-2,4-dienoic acid | 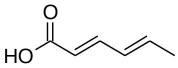 | + |
| **11** | Caproic acid | 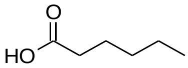 | + |
| **12** | Heptanoic acid | 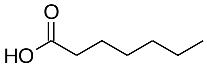 | + |
| **13** | Decanoic acid | 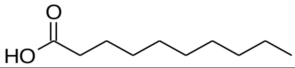 | + |
| **14** | Palmitic acid | 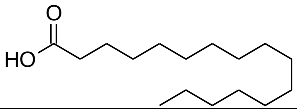 | + |

#### Supporting Figures

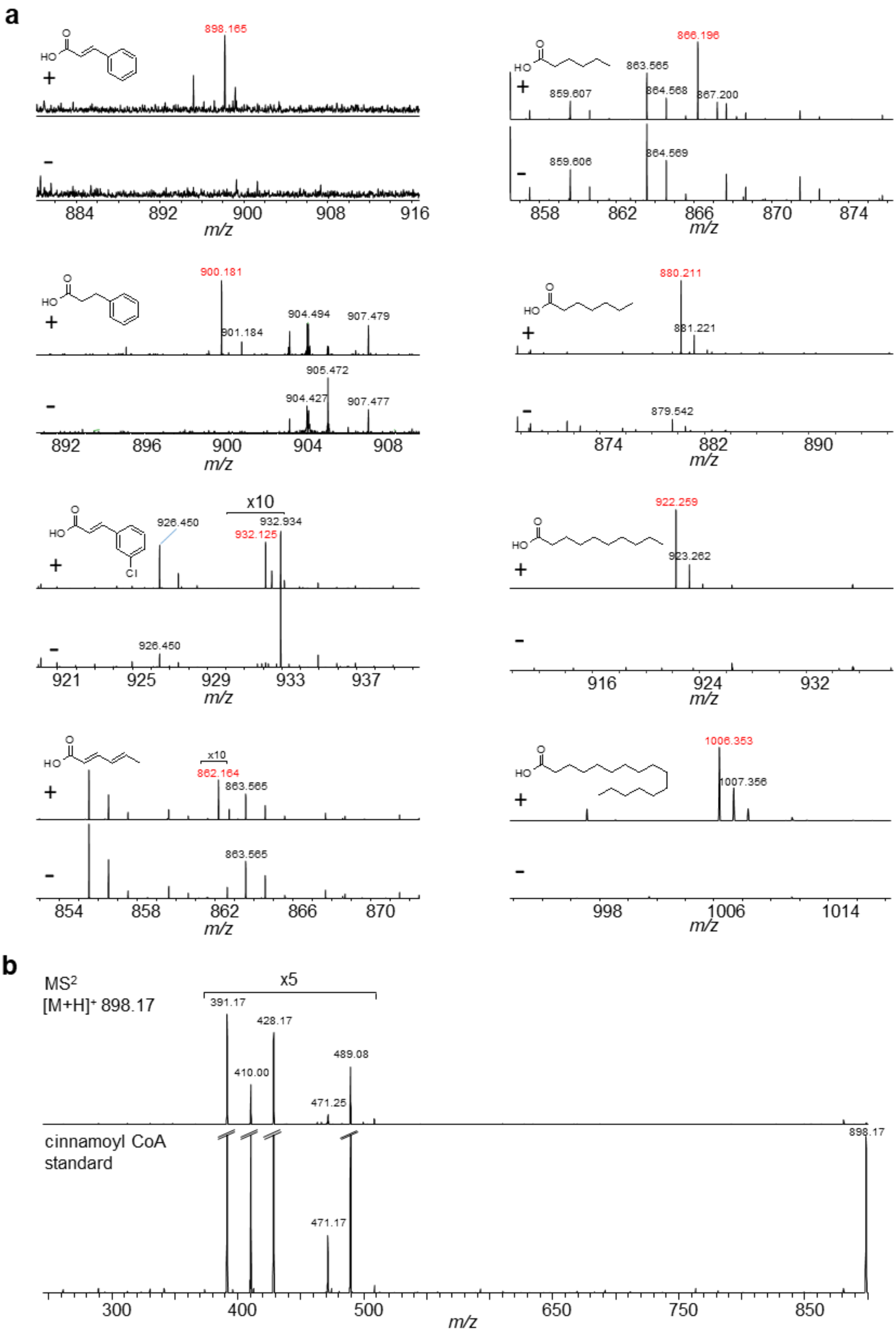

**Figure S1.** Detection of CoA-derivatives built upon the CoA-ligase activity of StlB. CoA ligase reaction was tested for substrates **1-14** (Table 1), CoA-esters were detected by *high resolution* MALDI-MS (**a**)**.** Due to the weak signal of cinnamoyl-CoA in MS$^1$, a further MS$^2$-experiment was performed in comparison with a cinnamoyl-CoA standard (**b**)**.**

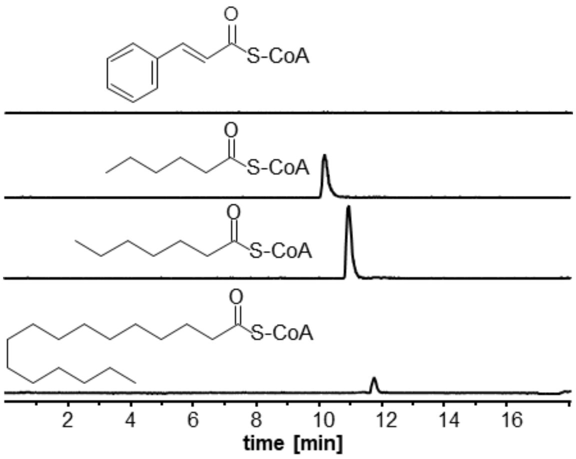

**Figure S2.** CoA-ligase reaction of StlB. Caproic ($C_6$)-, heptanoic ($C_7$)- and palmitic ($C_{16}$) acid were converted to their corresponding CoA ester by the CoA ligase activity of StlB. The conversion of cinnamic acid to cinnamoyl CoA was not detectable in this setup.

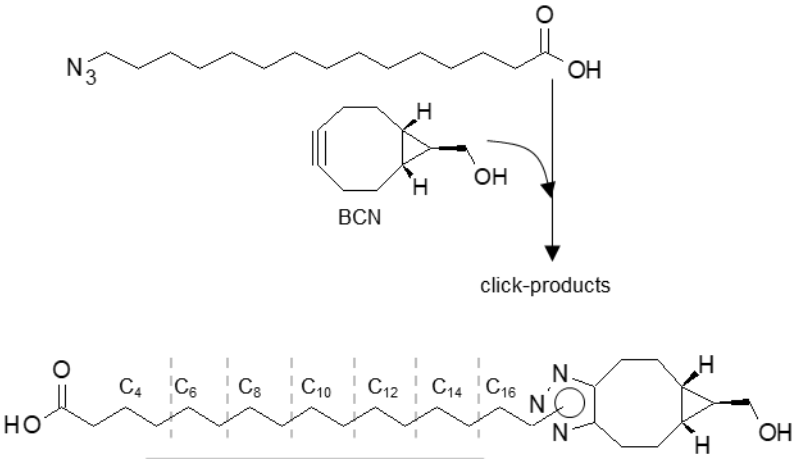

| Click product | calc. $[M+H]^+$ |
|---|---|
| **$C_{16}$** | 448.1 |
| **$C_{14}$** | 420.1 |
| **$C_{12}$** | 392.1 |
| **$C_{10}$** | 364.1 |
| **$C_8$** | 336.1 |
| **$C_6$** | 308.1 |
| **$C_4$** | 280.1 |

**Figure S3.** Expected click-products (table) after click-reaction with azido-palmitic acid and BCN.[3] HPLC-data, see Fig. S4

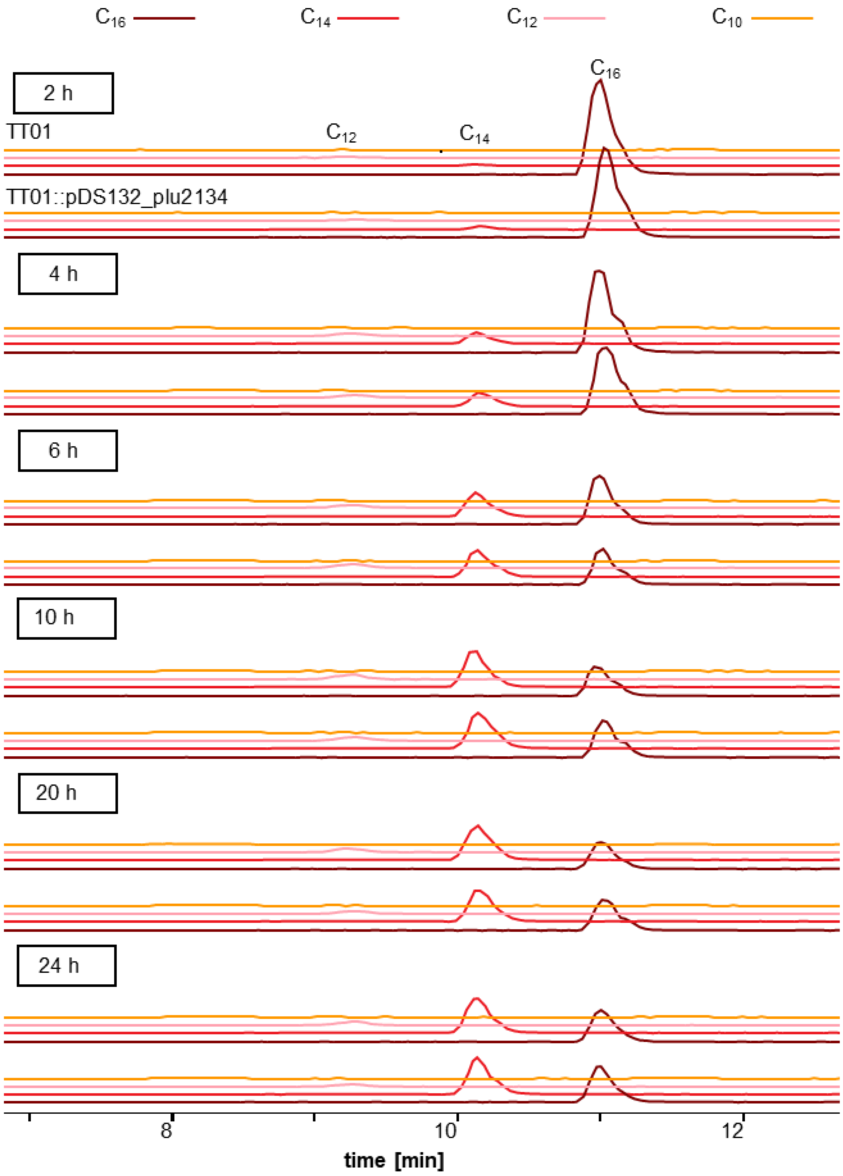

**Figure S4.** Fatty acid degradation profile of *P. laumondii* TT01 *stlB* (*plu2134*) plasmid insertion mutant compared to the wildtype. TT01 was fed with azido-labelled $C_{16}$-fatty acid and a click-reaction with BCN was performed to detect the degradation products ($C_{14}$, $C_{12}$, $C_{10}$, $C_8$) after 2 h, 4 h, 6 h, 10 h, 20 h and 24 h.

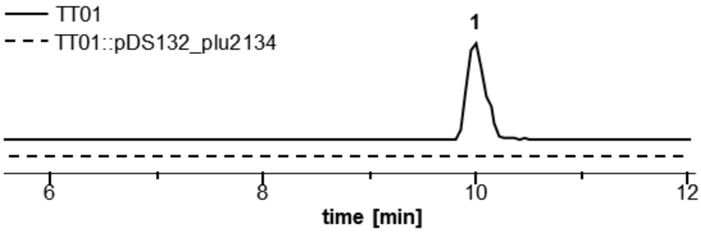

**Figure S5.** Growth of *P. laumondii* TT01 and insertion mutant TT01::pDS132_plu2134 (*stlB*) without antibiotic. Plasmid insertion of TT01::pDS132_plu2134 was verified by the loss of **1** production. **1** in TT01 was detected via HPLC-UV-MS at 9.9 min with a mass of 255.1 $[M+H]^{+\cdot}$ and its characteristic UV.

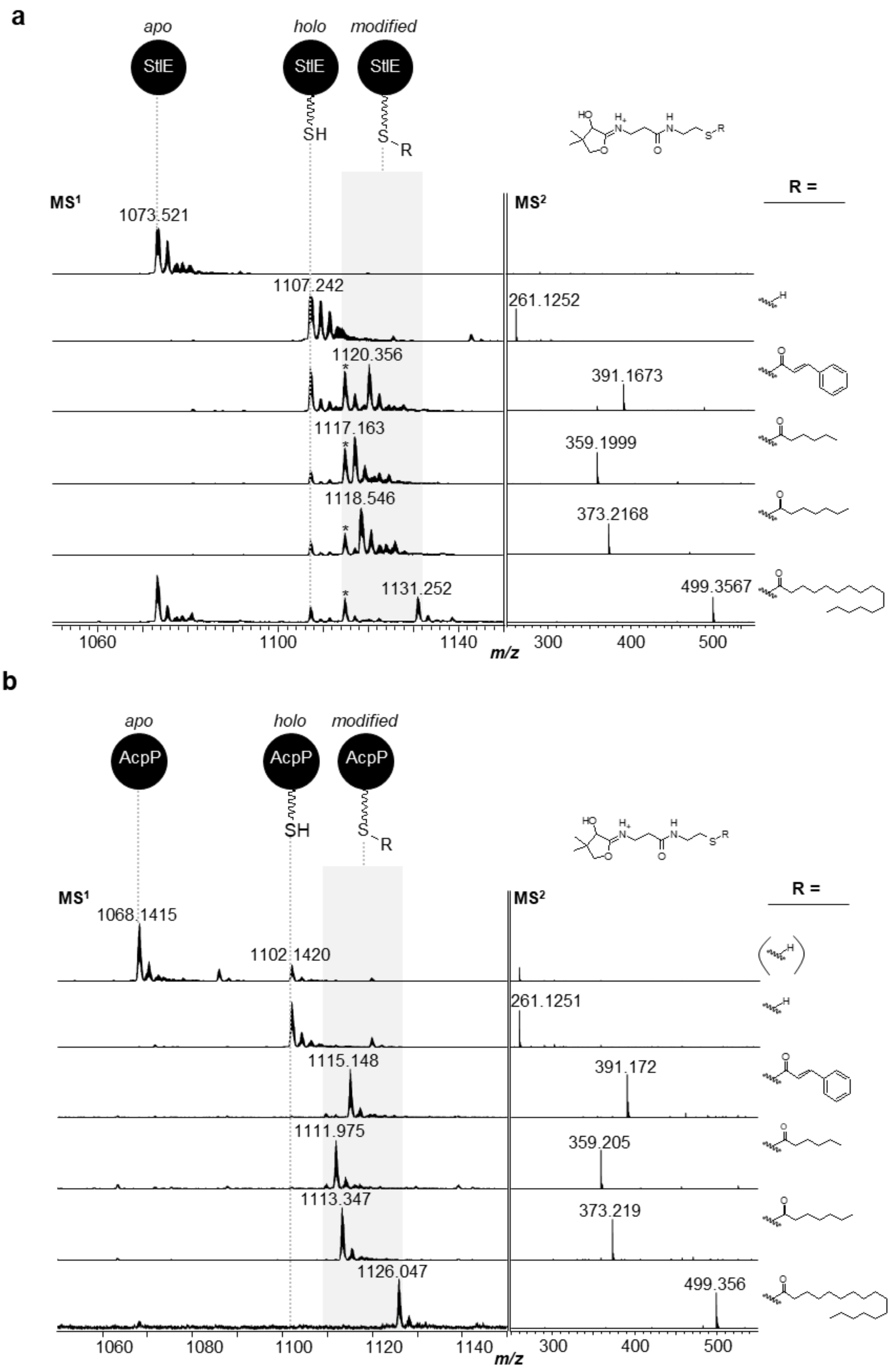

**Figure S6.** Loading of StlE (**a**) and AcpP (**b**) by AasS reaction of StlB with cinnamic, caproic, heptanoic and palmitic acid. The ACPs StlE and AcpP were transferred to their *holo*-forms by means of Sfp. In MS$^1$, the 10$^+$ charge state of the ACPs is shown and in MS$^2$, the corresponding Ppant ejection ions are displayed. In the case of StlE loading reactions, a side product was detected during overnight reaction (corresponding signals are marked with an asterisk).

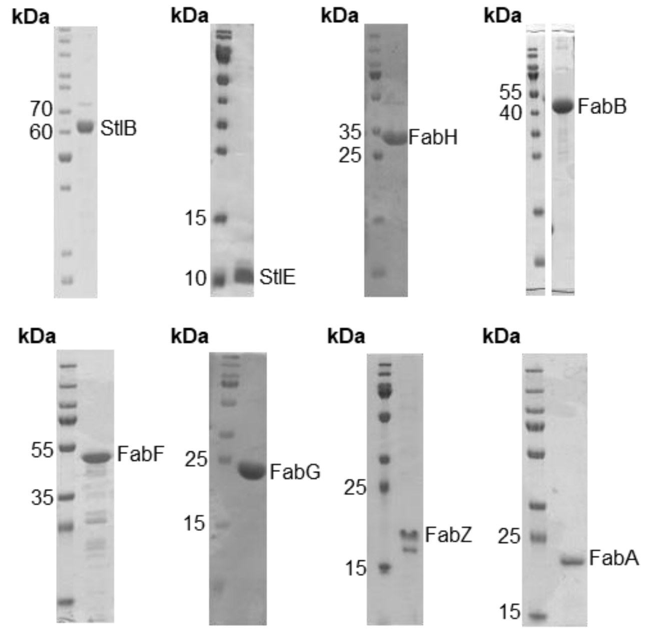

**Figure S7.** SDS-PAGE of strep-tagged StlB, StlE and FabH, FabB, FabF, FabG, FabZ, FabA.

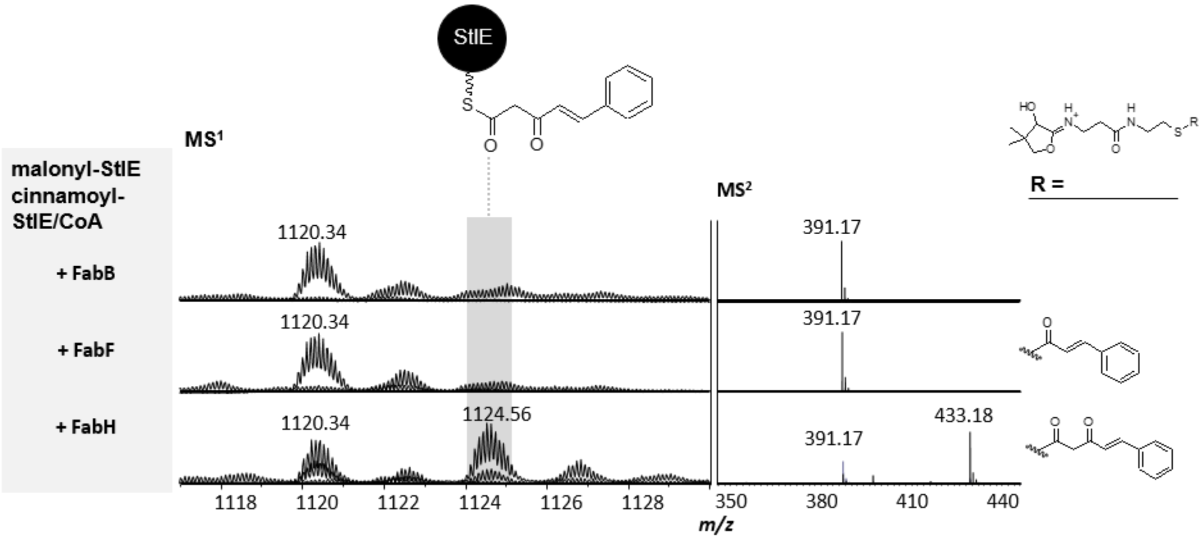

**Figure S8.** *In vitro* elongation with FAS II enzymes FabH, FabB and FabF and cinnamoyl-ACP as starter and malonyl-ACP as elongation unit. Spectra where overlaid as described in the experimental section.

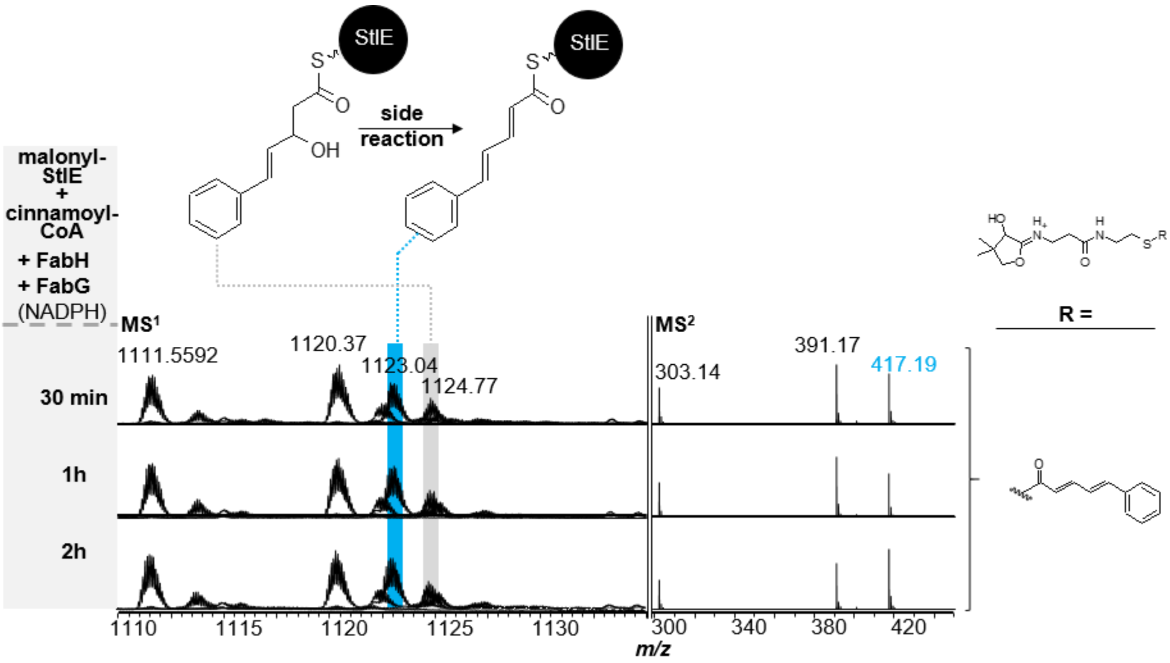

**Figure S9.** Time dependent *in vitro* analysis of side reaction with cinnamoyl-CoA as starter and malonyl-ACP as elongation unit being elongated and reduced by FabH and FabG (+NADPH), respectively. Spectra were overlaid as described in the experimental section. The dehydrated side product did not increase with increased reaction time.
